# Population Health Genetic Screening for Tier 1 Inherited Diseases in Northern Nevada: 90% of At-Risk Carriers are Missed

**DOI:** 10.1101/650549

**Authors:** JJ Grzymski, G Elhanan, E Smith, C Rowan, N Slotnick, S Dabe, K Schlauch, R Read, WJ Metcalf, B Lipp, H Reed, E Levin, J Kao, M Rashkin, J Bowes, A Bolze, K Dunaway, N Washington, A Slonim, JT Lu

## Abstract

In an unselected population of 23,713 participants who underwent clinical exome sequencing as a part of the Healthy Nevada Project (HNP) in Northern Nevada (Renown Health, Reno, Nevada) from March 15, 2018, to Sept 30, 2018 (Table S1) we find a 1.26% carrier rate for expected pathogenic and likely pathogenic genetic variants in (FH: *LDLR, PCSK9, APOB*), Hereditary Breast and Ovarian Cancer (HBOC: *BRCA1, BRCA2*) and Lynch Syndrome (LS: *MLH1, MSH2, MSH6, PSM2*) with over 90% of carriers undetected under current medical practice. 26% of carriers were found to have advanced disease with 70% first diagnosed before the age of 65. Less than 20% of all carriers had any documented suspicion for inherited genetic disease in the medical record and upon direct follow-up survey under 40% of carriers had family history of relevant disease. A population preventative genetic screening approach for patients under 45 may improve outcomes.

## Main

Here we report on genetic risk and disease manifestation for the above three inherited autosomal dominant diseases (9 genes) that have evidence-based recommendations for genetic testing targeted towards subsets of the population based on family history, ethnic background, or other demographic characteristics ^1^. Interrogating the clinical genetic data for previously described pathogenic and likely pathogenic (P/LP) variants ^2^ identified 290 individuals for a population prevalence of 1.26% (Table S1). Although not corrected for familial relatedness or potential ascertainment bias, the prevalence rate of BRCA related HBOC (∼1 in 185), LS (∼1 in 520) and FH (∼1 in 208) are in line with recent reports ^3–5^. Carriers broadly resemble the overall demographic profile of the HNP with a median age of 47, 63% female, and 78% caucasian (Table S1). We then curated and reviewed available medical records, extracted and processed in February 2019, and combined this data with surveyed family history disease risk (See Online Methods) for each of the 243 participants (83.3% of the 290 participants) who are patients at Renown Health System. We find that 62 (26%) patients had already presented with disease. First disease diagnosis was ascertainable for only 55 patients (of the 64) as some patients enter Renown Health as patients with disease. The median age of first diagnosis for these patients was 60 with 88.6% of disease first presenting after the age of 40 (Figure 1), and 70% before the age of 65. Of these patients, 26 (47.2%) had no documented diagnostic information that could have raised suspicion of underlying genetic condition prior to diagnosis, suggesting a somewhat acute presentation.

**Figure 1:**
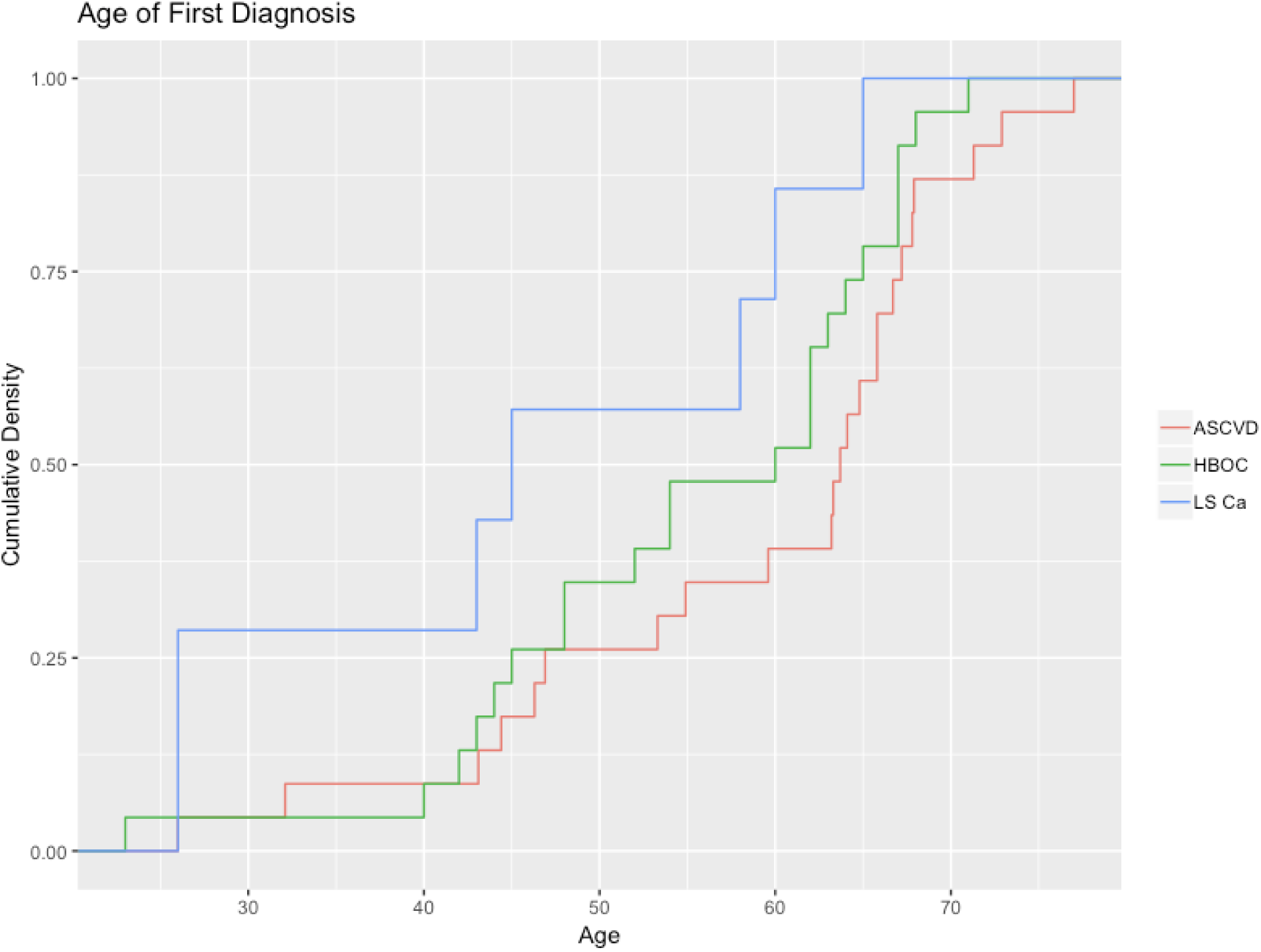
Empirical cumulative distribution for age of first diagnosis for FH related Atherosclerotic cardiovascular disease (ASCVD), BRCA1/2 related Hereditary Breast and Ovarian Cancer (HBOC) and Lynch Syndrome related malignancy (LS Ca).

For HBOC carriers, we were able to review 108 medical records. These patients had a median age of 51 with 91 (84.3%) of the carriers falling outside of NCCN practice guidelines for genetic testing. We also find that 23 of the HBOC carriers (21.2%) had a diagnosis of breast or ovarian cancer with a median diagnosis age of 60. In comparison, the prevalence of non-BRCA1/2 HBOC in women at age 55 is ∼1.47%^6^. 9 of the 23 diseased carriers (39.1%) were within NCCN guidelines for genetic testing; however, only five of the nine had genetic testing prior to HNP, likely due to early onset of disease (median age at diagnosis of 45 for tested individuals). The remaining four had a median age of diagnosis of 64.5, which likely reduced clinical suspicion of germline disease. We also found that 14 (60.9%) of the carriers with disease were nonetheless tested after disease diagnosis even though they fell outside of NCCN guidelines (Table 1). Although we find no differences in disease-free progression between BRCA1 and BRCA2 carriers (Figure S1) in our cohort but do find that family history provides incremental risk excess of carrier status alone (Figure S2). This is consistent with previous studies that show that family history is an independent risk factor for breast/ovarian cancer^7^. Lastly we find that for 15 of the 23 (65.2%) of the diseased patients, there was no diagnostic evidence that could indicate suspicion for disease prior to diagnosis with breast or ovarian cancer. Instead, these individuals either presented acutely or through routine clinical screening for breast cancer (i.e. mammography).

**Table 1:** Detailed information about HBOC, LS, and FH including carriers per gene and disease prevalence

|  | HBOC |  |  |  | LS |  |  |  |  |  | FH |  |  |  |
| --- | --- | --- | --- | --- | --- | --- | --- | --- | --- | --- | --- | --- | --- | --- |
|  | BRCA1 | BRCA2 | Total |  | MLH1 | MSH2 | MSH6 | PSM2 | Total |  | APOB | LDLR | PCKS9 | Total |
| # of Carriers | 51 | 84 | 135 | # of Carriers | 8 | 4 | 17 | 17 | 46 | # of Carriers | 26 | 79 | 12 | 117 |
| # w/ Medical Records | 42 | 66 | 108 | # w/ Medical Records | 5 | 3 | 16 | 13 | 37 | # w/ Medical Records | 20 | 67 | 11 | 98 |
| Sex |  |  |  | Sex |  |  |  |  |  | Sex |  |  |  |  |
| Female | 26 | 36 | 62 | Female | 5 | 1 | 10 | 8 | 24 | Female | 13 | 46 | 6 | 65 |
| Male | 16 | 30 | 46 | Male | 0 | 2 | 6 | 5 | 13 | Male | 7 | 21 | 5 | 33 |
| Median Age [Range] | 45 [18-81] | 52 [19-81] | 51 [18-81] | Median Age [Range] | 51 [26-66] | 28 [28-31] | 57 [26-84] | 68 [26-84] | 58 [26-84] | Median Age [Range] | 46 [19-79] | 47 [18-77] | 56 [27-81] | 47 [18-81] |
| # of Patients with indication for genetic testing | 17 |  |  | # of Patients with indication for genetic testing | 2 |  |  |  |  | # of Patients with indication for genetic testing | 3 |  |  |  |
| # w/ Genetic Testing Completed (July 2018) | 8 (47.9%) |  |  | # w/ Genetic Testing Completed (July 2018) | 2 (100%) |  |  |  |  | # w/ Genetic Testing Completed (July 2018) | 2 (67%) |  |  |  |
| # of Patients without indication for genetic testing | 91 |  |  | # of Patients without indication for genetic testing | 35 |  |  |  |  | # of Patients without indication for genetic testing | 95 |  |  |  |
| # w/ Genetic Testing Completed (July 2018) | 5 |  |  | # w/ Genetic Testing Completed (July 2018) | 1 |  |  |  |  | # w/ Genetic Testing Completed (July 2018) | 2 |  |  |  |
| # with breast/ovarian CA | 10 | 13 | 23 | # w/ LS related malignancy | 3 | 3 | 6 | 3 | 15 | # with ASVCD | 6 | 18 | 2 | 26 |
| Sex |  |  |  | Sex |  |  |  |  |  | Sex |  |  |  |  |
| Female | 9 | 13 | 22 | Female | 3 | 1 | 4 | 3 | 11 | Female | 4 | 8 | 0 | 12 |
| Male | 1 | 0 | 1 | Male | 0 | 2 | 2 | 0 | 4 | Male | 2 | 10 | 2 | 14 |
| Median age [Range] at time of diagnosis | 62.5 [40-67] | 52 [23-71] | 60 [23-71] | Median age [Range] at time of diagnosis | 45 [26-65] |  |  |  |  | Median age [Range] at time of diagnosis | 53 [32-72] | 65 [32-70] | 57 | 62 [32-72] |
| Prior Clinical Suspicion of Genetic Disease | 3 | 5 | 8 | Prior Clinical Suspicion of Genetic Disease | 1 | 2 | 2 | 0 | 5 | Prior Clinical Suspicion of Genetic Disease | 5 | 6 | 1 | 12 |

For LS carriers, 37 medical records were available and reviewed. These patients had a median age of 58 with 35 (94.6%) of the carriers falling outside of NCCN practice guidelines (Table 1) for genetic testing. Carriers of LS P/LP variants had marked increases in oncologic events; 13 of the 37 (35.1%) carriers had a colon, ovarian or other cancer diagnosis with a median age of first diagnosis of 44. Seven of the 13 had colon cancer while six of the 13 had another type of malignancy; 4 patients presented with malignancy in two or more locations. In contrast the prevalence of colon cancer at ages 40-49 is 0.12%^6^ (Table 1) in individuals without LS. The prevalence rate in our unselected population is in-line with the approximately ∼40% cumulative incidence rates of cancer by age 50 in family history-selected LS mutation carriers ^8^. Of the seven patients with colon cancer, six had documentary evidence in the medical record that could have raised suspicion for an underlying genetic condition, however only 2 of these patients fell within NCCN guidelines and were tested for Lynch Syndrome. No other patients had genetic testing. This is perhaps unsurprising given that average risk CRC screening starts at age 50^9^. In contrast, annual screening in LS patients is recommended at age 20^10^.

In FH carriers, 98 medical records were reviewed along with family history ascertained during the Return of Results (See Online Methods). We found that 63% of FH carriers did not have known family history of heart disease; in addition, only three individuals were found to have an indication for genetic testing and only two had testing. The remaining 95 (96.9%) of carriers were found to be outside of guidelines-based testing^11^. Of the 98 carriers, 26 (26.5%) had manifested with atherosclerotic cardiovascular disease (ASCVD). We found a relative enrichment of 2.3 (95% CI: 1.13-3.47, Two Proportion Test) of males to females; 14 of 33 (42.4%) of male carriers (median age of diagnosis 64) and 12 of 65 (18.5%) of female carriers (median age of diagnosis 63) had manifested ASCVD (Table 1). This is in line with previously described rates for phenotypic FH ^12^. In comparison, the prevalence for non-FH ASCVD in 40-59 y.o males and females is 6.3% and 5.6% ^13^. We also observed no discernable differences in disease penetrance between APOB, PCSK9 and LDLR (Figure S3). Lastly, there were 8 major acute events out of the 26 disease cases: five myocardial Infarctions (MI) and three cerebral vascular accidents (CVA). For six of these eight events, the first documented ASCVD related event was the acute event. The remaining patients all had evidence of coronary intervention.

To our knowledge, the ascertainment of genetic disease in an unselected population using a population health approach has not previously been performed. While the observed prevalence rates in HNP for carriers of FH, HBOC, and LS are consistent with previously published biobanking studies ^3–5^, we discover that current medical guidelines are largely inadequate for detecting carriers of these three diseases. Over 90% of P/LP carriers are undetected today because guidelines are narrowly focused on selected populations. Yet disease penetrance for LS and FH in our unselected population is similar to that in selected populations recommended by guidelines, and substantially elevated in HBOC carriers (albeit at a lower rate than those with additional family history). Our study also illustrates that average risk screening protocols may not adequately identify at-risk individuals as average risk screening typically is less intensive and starts later in life: screening using mammography for breast cancer starts at age 50 in women ^14^, for colon cancer at age 50 in both sexes^15^, and for lipid disorders at age 35 and 45 for men and women^16^, respectively.

The HNP follows a public health community-based approach with direct participant outreach, enrollment and return of results. This also includes direct to participant genetic education and physician CME courses (See Online Methods). Some self-selection for the study is evident from the relative enrichment of caucasians and the higher educational attainment of the cohort. However these demographics typically enrich for higher levels of engagement with health systems^17^ and higher compliance with preventative care. Our analysis of medical records is also limited to 12 years of electronic health records from the Renown Health System, although this is the dominant health system in Northern Nevada with more than 70% market share.

In the over 1.1 million health records at Renown Health, diagnosis of FH, LS or HBOC occurs in less than 0.2% of patients. Our results suggest that genetic predisposition testing as a first line preventative genetic screen for these conditions could lower mortality and morbidity related to inherited disease risks that are today insufficiently detected under current medical practice. A population health screening approach to genetic medicine requires different scrutiny given the potential for added costs, over-interpretation of disease risk, and ethical and social factors^18^. Cost-effectiveness studies have found predispositional genetic screening with return of a limited panel of genetic information to be cost effective in a population under 45 years old when testing costs are in the low hundreds of dollars^19,20^. Our experience with the Healthy Nevada Project suggests that a public health based approach to genetic screening in younger patients can provide substantial clinical, economic, and patient benefit.

**Figure S1:**
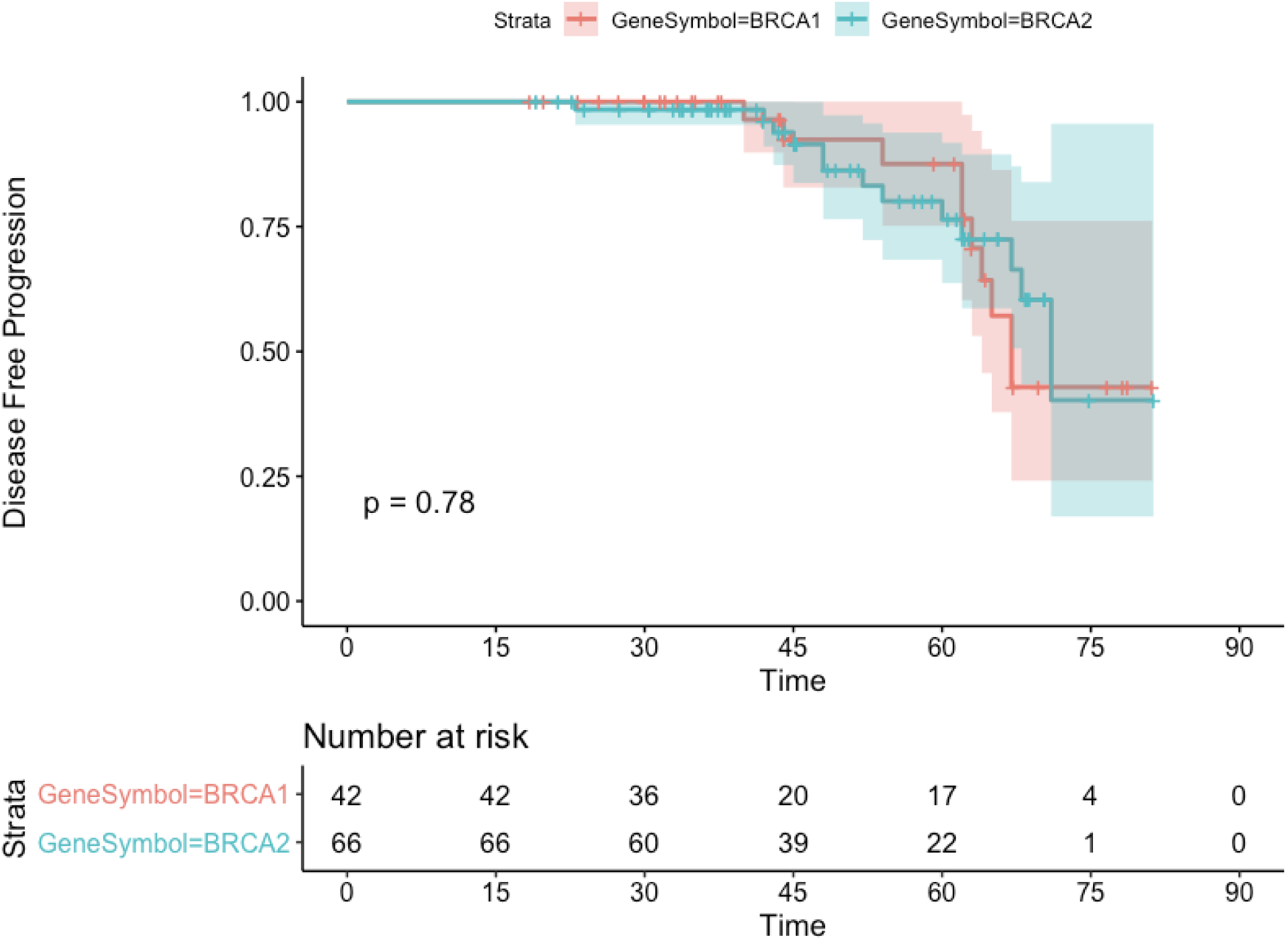
No difference in disease free progression between carriers of BRCA1 and BRCA2 P/LP variants.

**Figure S2:**
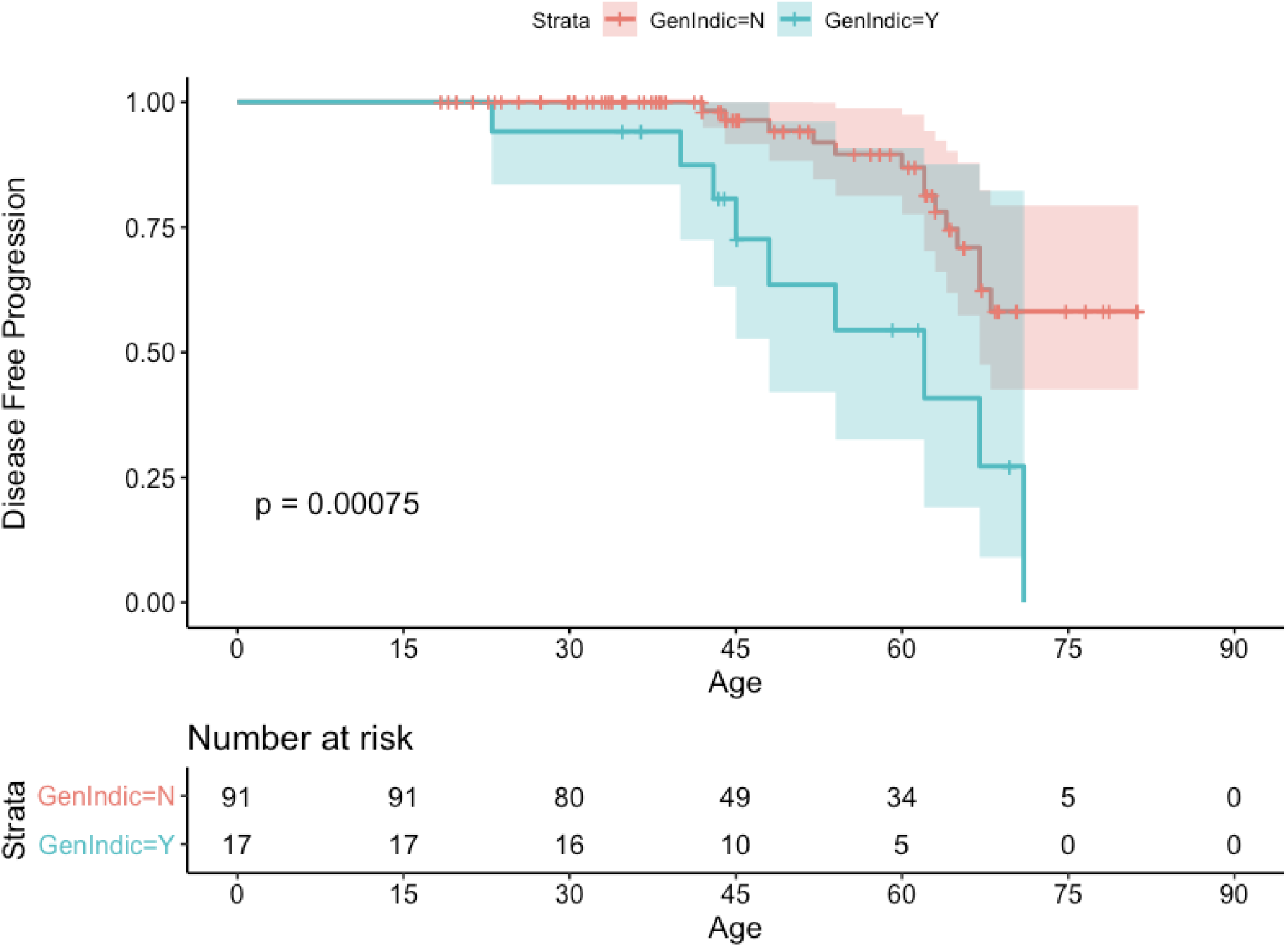
Disease-free progression in HBOC carriers that are within and outside of NCCN guidelines for testing.

**Figure S3:**
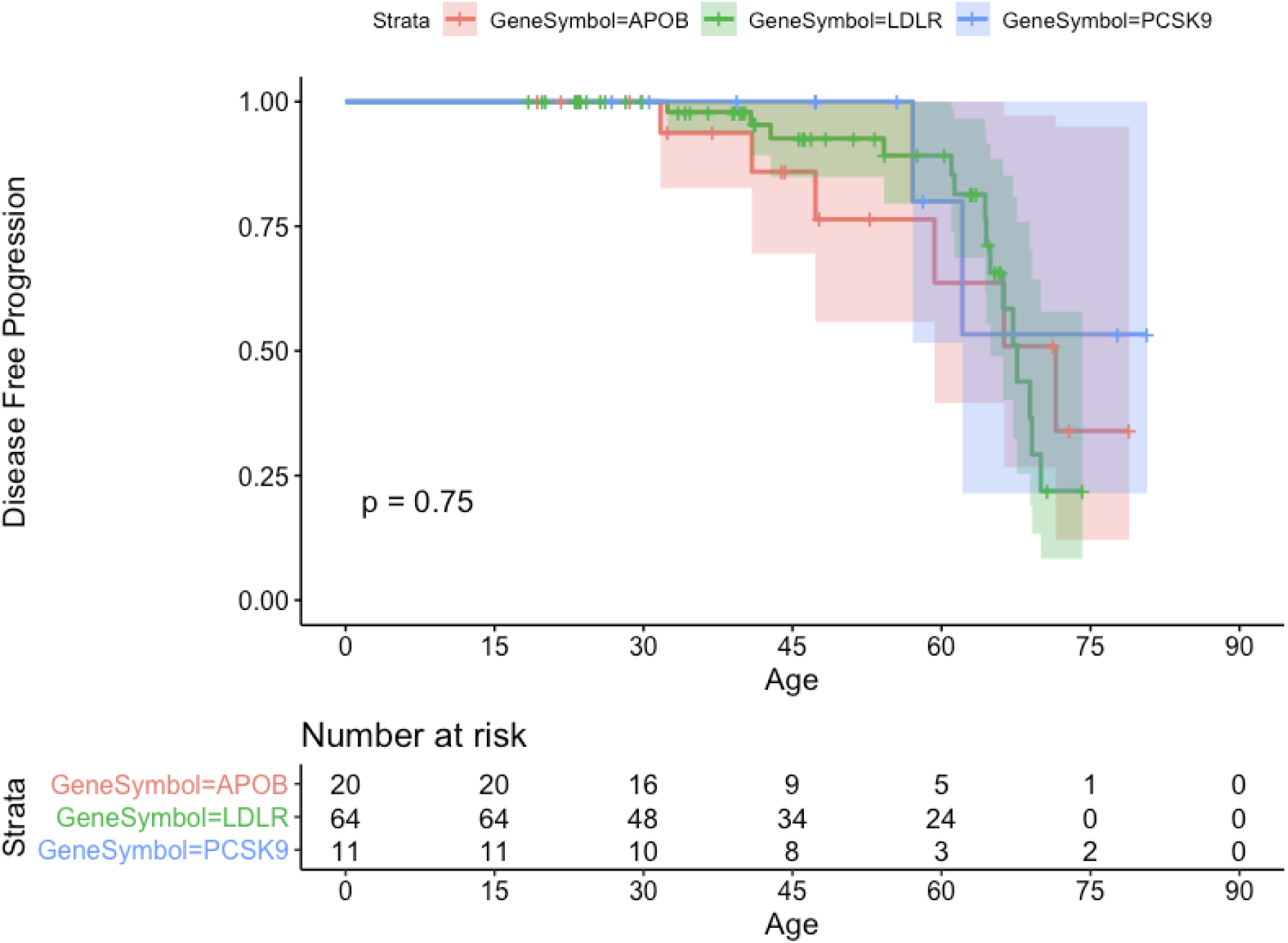
No difference in disease free progression between carriers of APOB, LDLR, and PCSK9.

**Table S1:** Demographic Profile of Healthy Nevada Project Participants.

| **Race/Ethnicity:** | **HBOC carriers** | **FH Carriers** | **LS Carriers** | **Healthy Nevada Project** | **Renown Health System** | **Washoe County** |
| --- | --- | --- | --- | --- | --- | --- |
| Count | 135 | 117 | 46 | 23,709 | ~1.03M | 421,407 |
| Prevalence | 0.57% | 0.49% | 0.19% |  |  |  |
| Sex |  |  |  |  |  |  |
| Male (%) | 39% | 35% | 33% | 34% | 51% | 50% |
| Female (%) | 61% | 65% | 67% | 66% | 49% | 50% |
| Age (Median, Stdev) | 44 [16.9] | 47.2 [18.2] | 50.5 [19] | 44.5 [19] | 49 [24] | 38 |
| Ethnicity (Count, %) |  |  |  |  |  |  |
| Caucasian | 107 (79.3%) | 87 (74.4%) | 38 (82.6%) | 81% | 72% | 63% |
| Hispanic/Latino | 17 (12.6%) | 6 (5.1%) | 1 (2.2%) | 10% | 10% | 25% |
| Asian | 4 (3%) | 11 (9.4%) | 3 (6.5%) | 3% | 3% | 5% |
| Native American | 3 (2.2%) | 0 (0%) | 2 (4.3%) | 1% | 1% | 1% |
| Pacific Islander | 0 (0%) | 0 (0%) | 0 (0%) | 1% | 0% | 1% |
| African American | 2 (1.5%) | 8 (6.8%) | 1 (2.2%) | 1% | 3% | 2% |
| Other/Unknown | 2 (1.5%) | 5 (4.3%) | 1 (2.2%) | 3% | 11% | 3% |

**Table S2:** List of P/LP Variants for all cases

| Disease | Gene | # of Cases | Relative Contribution | Population Prevalence |
| --- | --- | --- | --- | --- |
| HBOC | BRCA1 | 51 | 37.78% | 0.215% |
| HBOC | BRCA2 | 84 | 62.22% | 0.354% |
| LS | MLH1 | 8 | 17.39% | 0.034% |
| LS | MSH2 | 4 | 8.70% | 0.017% |
| LS | MSH6 | 17 | 36.96% | 0.072% |
| LS | PMS2 | 17 | 36.96% | 0.072% |
| FH | APOB | 26 | 22.22% | 0.110% |
| FH | LDLR | 79 | 67.52% | 0.333% |
| FH | PCKS9 | 12 | 10.26% | 0.051% |

## Online Methods

### Healthy Nevada Project

The Healthy Nevada Project (https://healthynv.org/) is an all-comers human subject research study on health determinants in Northern Nevada with specific recruitment foci in rural and socio-economically depressed areas in Northern Nevada. This ongoing program, with a targeted enrollment of 250,000, targets a diverse demographic: 23.8% of individuals are recruited from rural zip codes and ∼20% of the study consists of minority participants, predominantly hispanic and native american (Table S1). Participants consent to a research protocol that includes (1) clinical “Exome+” sequencing, (2) linking of sequencing data with electronic health records, (3) return of secondary findings and (4) recontact for health surveys and participation in future studies. 99.2% agreed to return of secondary findings and 96.1% agreed to recontact for future studies. The project 1106618-15, was approved by the University of Nevada, Reno Institutional Review Board.

### Return of Secondary Findings

The return of secondary findings process started in October 2018 and was completed for these patients by February 2019. For each positive secondary finding the de-identified participant ID was cross-matched with a de-identified consent. If the participant consented to return of results, the data was re-identified by an authorized member of the research team. Contact information was then linked by a designated research coordinator at the Office of Research and Education using Renown Health’s secure email platform. Linked contact list and individual reports were stored in a secure folder on an encrypted server only accessible to designated research staff.

Clinical Research Coordinators scheduled a 15 minute phone appointment to speak to a clinician regarding their results. Participants were called 1-2 days prior to the first available appointment to minimize wait time for the delivery of their results over the phone. If there was no answer, a voicemail was left with the participant. A phone call was also made to any secondary number on file, and a message left if applicable. The voicemail was then followed up with an email to the same participant with similar information to call back and schedule an appointment. The date of scheduling or unsuccessful attempts to contact each participant was recorded on the contact list. Every effort was made to minimize the significance of the call to the greatest extent possible until the day of the scheduled appointment to avoid undue stress to the participant.

All medically significant results were delivered by a clinical provider with appropriate credentialing to deliver genetic results. To ensure consistency between participants, a generalized script was used by the clinician when delivering results. At the time of the appointment, the clinician verified that the participant still wanted to be informed of any findings that might be relevant to their health prior to continuing the call. Once confirmed, the clinician informed the individual of their secondary finding, what it meant for their health, and the importance of following up with a medical professional upon receiving their results. Family history pertaining to their results was collected for research purposes using Renown’s family history questionnaire. This questionnaire was designed to survey family medical history specific to (1) NCCN guidelines for Lynch Syndrome, (2) Tyrer-Cuzick ^21^, or Penn II empirical risk calculators ^22^ for HBOC, and (3) family disease risk for cardiovascular disease ^12,23^. Prior to finishing the call, participants were asked if they would like a copy of their results delivered to them either by secure email and/or through the postal service.

Once a participant was notified of their results, a copy of their signed report was sent to their preferred method of delivery (email or letter), along with supporting documents. Supporting documentation included: a cover letter with contact information for the study, templated letters they can provide to their physicians and family members, a copy of available primary care physicians in the area if requested by the participant, and any other educational or informational material pertaining to their findings as deemed applicable. Source documents for the return of results were maintained by research staff per research policy. In addition, a copy of returned results is retained in our Clinical Trial Management System for compliance with study protocols.

### Electronic health record analysis

HNP participant IDs were mapped to medical record IDs. For all HNP participants with valid medical record IDs, clinical data were retrieved from the Renown Epic Clarity database (February 2019) utilizing a script. The script was designed to collect patient diagnoses from multiple Clarity tables including, but not limited to, encounter diagnoses, problem lists, medical histories and claims, and invoices. The script also de-identified records by removing all PHI. The diagnoses retrieved were encoded in Renown’s proprietary codes and were mapped to ICD10-CM and ICD9-CM diagnosis codes utilizing a mapping schema. For each patient, each unique ICD diagnosis code was tagged with the date of its first and last occurrence across all the Clarity tables that were queried by the script. Data regarding lipid profile results and evidence of prescriptions of lipid lowering therapies and their types were retrieved in a similar manner.

For each patient, diagnosis codes, lab data and medication orders were reviewed by a clinician. Diagnosis codes that were not deemed relevant to the underlying FH, LS and HBOC conditions were removed. The remaining codes, laboratory tests, medication orders and timestamps were used to determine the status of the condition itself, family history and other risk factors. This includes determining if 1) the patient had genetic testing prior to Return of Results from the HNP, 2) if the patient’s family history and patient conditions supported clinical suspicion for an underlying genetic condition and 3) if the patient had a diagnosis of a disease that could reasonably be associated with underlying genetic risk such as hereditary breast & hereditary ovarian cancer, Lynch Syndrome associated malignancies and /or ASCVD related diagnoses and the earliest date of diagnosis.

### Assay and Sequencing

Sequencing for all participants was performed at Helix’s CLIA (#05D2117342) and CAP Accredited (#9382893) facility in San Diego, CA. The Exome+ (Helix) is a proprietary exome that combines a highly performant medical exome with a microarray backbone into a single sequencing assay (www.helix.com). The Exome+ provides highly uniform and complete coverage across medical regions of the genome, with a target coverage >20x for >99% of all targeted bases in this study, for all participants. All returned variants meet Helix’s validation criteria for clinical return. Performance specification can be found in Helix’s Exome+ Performance White Paper (https://cdn.helix.com/wp-content/uploads/2017/07/Helix-Personal-Genomics-Platform-White-Paper.pdf).

### Selection of Pathogenic/ Likely Pathogenic Alleles

The list of pathogenic/ likely pathogenic were filtered from CLINVAR using filters for pathogenic or likely pathogenic variants. We then removed any variants with conflicting interpretations and any variants whose origin was somatic. These data were generated from the June 3, 2018 CLINVAR database (https://www.ncbi.nlm.nih.gov/clinvar/).

## Data Availability Statement

The data that support the findings of this study are available from the corresponding author upon reasonable request.

## Code Availability Statement

The code that supports the findings of this study are available from the corresponding author upon reasonable request.

